# Fecal and skin microbiota of two rescued Mediterranean monk seal pups during rehabilitation

**DOI:** 10.1101/2023.07.05.546574

**Authors:** Aggeliki Dosi, Alexandra Meziti, Eleni Tounta, Kimon Koemtzopoulos, Anastasia Komnenou, Panagiotis Dendrinos, Konstantinos Kormas

**Affiliations:** Department of Ichthyology and Aquatic Environment, University of Thessaly, 384 46 Volos, Greece; MOm/Hellenic Society for the Study and Protection of the Monk Seal, Solomou 18, 10682, Athens, Greece; School of Veterinary Medicine, Faculty of Health Sciences, Aristotle University of Thessaloniki, 574 00 Thessaloniki, Greece

**Keywords:** conservation, marine mammal, red list, threatened, *Monachus monachus*, host-microbe interaction

## Abstract

The role of animal host-associated microbiomes is becoming more apparent and defined for wild animals, especially for the species under conservation strategies. This study investigated the succession of fecal and skin bacterial microbiota of two rescued female Mediterranean monk seal (*Monachus monachus*) pups for most of their rehabilitation period. Bacterial species richness and diversity was assessed by high-throughput sequencing of nine freshly collected fecal samples and four skin swabs per individual. Both the fecal and skin microbiota highly overlapped in their containing operational taxonomic units (OTUs) and abundance patterns. The fecal microbiota was separated in two distinct periods, and was dominated by OTUs related to the *Shigella*, *Streptococcus*, *Enterococcus*, *Lactobacillus* and *Escherichia* genera in the first period, while in the second period the dominating genera were the *Clostridium, Blautia, Fusobacterium, Edwardsiella* and Bacteroides. The skin microbiota was highly similar between the two individuals in each sampling and were dominated by *Psychrobacter-, Elizabethkingia-* and *Bergeyella*-related OTUs. The provided antibiotic treatment along with the provided probiotics and nutritional supplements, resulted in a major turnover of the bacterial microbiota with the potentially detrimental OTUs being eliminated towards the end of the rehabilitation period, prior to the release of the pups in the wild.

## INTRODUCTION

The introduction of the holobiont and hologenome concepts (1), intrigued the scientific interest on the investigation of the associations and interactions between wild animals and their microorganisms (2, 3). This research promotes the notion that the animals as hosts have inseparable evolutionary and functional roles with their microbiomes (4). Recently, the concept that our view on animals should shift from the object (animal organism) to the process (animal organisms + its everchanging microbiome) ontology has been proposed (5). Such tight and vital symbiotic relationships between macroorganisms and their microorganisms are pivotal for the host’s health, development, and nutrition and for this there is no reason not to consider them important, especially during ecological disturbances Host-microbe interactions work, among other functions, as buffers against various kinds of biological or environmental disturbances to maintain the host’s homoeostasis and function Indeed, similar rescuing roles have been proposed for environmental microbiomes, too (8).

It is recently estimated that 42,100 plant and animal species face extinction, with 27% of them being mammals (9). This sets the need for intensifying our current and effective conservation strategies but also to search for more novel approaches. The development of such strategies requires a more holistic knowledge of the organisms under threat. Soon after the first case studies showing the importance of microbiomes in conservation issues of wild animals (e.g., (10, 11)) the conceptual importance of animal microbiomes was revealed (12, 13). The field has progressed so fast that even well-established human microbiome-based manipulations for therapeutic purposes are now applied for captive animals to improve the animals’ health (14). Regarding animal species with both natural (wild) and captive (zoos, laboratory animals, farmed species, etc.) populations the comparison between the two types of microbiomes is considered as the first important step towards knowing the animals natural microbiome and/or the impact of biological or environmental disturbances (15–18). More specifically, for endangered animals which are under targeted protection actions, microbiome analysis is often restricted to only captive animals’ hospitalization or rehabilitation of individuals from the wild for enhancing the species natural population.

Despite that the total biomass of marine mammals across the world’s oceans is only 0.3% of the total marine animal biomass, when arthropods and fish have 38.6 and 27.0% (19), several of these species are endangered and protected due to anthropogenic activities, like the Mediterranean monk seal, *Monachus monachus* (20). More than half of the estimated total marine mammal biomass (ca. 40 Mt), is attributed to baleen whales, leaving seals with a low biomass contribution (21). However, apart from being the most endangered pinniped species in the world, *M. monachus*, the only species of the *Monachus* genus (22, 23), has several other points which attract the scientific interest. Although once abundant throughout the Mediterranean, in the Black Sea,, in the north-west Atlantic coast of Africa, the Canary islands, the Azores and the Madeira archipelago, its distribution is now limited to the Eastern Mediterranean Sea, an isolated population in the Atlantic coast of Africa (Mauritania) and a small isolated population in Madeira due to human-induced population declines though persecution and habitat destruction (24). Today, the species is designated as “Endangered under criteria D” by the International Union for Conservation of Nature (IUCN) List of Threatened Species (25) with signs of improving population through targeted scientific research, monitoring of local seal populations, education, public awareness and citizen science campaigns, and rescue and rehabilitation of wounded, sick, and orphaned seals (24, 26, 27). The latter, includes animal care practices on land for both young and adult individuals with specific medical and targeted nutritional care.

To date, there is no available scientific literature on any *M. monachus* microbiota or microbiome, despite that it has been shown that the Phocidae family seem to clearly differentiate in their fecal microbial communities compared to terrestrial carnivores (28). The only available relevant knowledge stems from its closely related species, the Hawaiian monk seal (*Neomonachus schauinslandi*) and refers to the identification of either bacterial antibodies of hauled out specimens (29), cultured aerobic bacteria of the upper respiratory tract of captive animals (30) and bacterial pathogen prevalence in the blood serum of experimentally resident and translocated seals (31). One reason for such restricted knowledge is related to the multiple challenges raised in microbiome sampling from marine mammals. However, several of these marine mammalian species are hospitalised, rehabilitated or live under captivity in zoos where their microbiomes, although far from that of their natural counterparts, are far more reachable. In this paper, we report for the first time on the fecal and skin microbiota succession of two rescued female *M. monachus* pups (Lena and Nicole) during their 5-month rehabilitation period prior to their release in the wild. We evaluated the impact of the provided medication and feeding scheme to these microbiota profiles by analyzing the changes of the skin and fecal bacterial community composition.

## RESULTS

After applying quality filtering and chimera removal of the 16S rRNA gene V3–V4 region amplicons, a total of 807,391 sequences were retrieved. The number of sequences per sample was rarefied to be equal to the smallest number (29,817) of sequences per sample. These sequences were assigned to 328 unique OTUs at a similarity cut-off level of 97%. The dominant bacterial phyla in the whole data set were Firmicutes, Gammproteobacteria, Bacteroidota, Fusobacteria and Actinobacteria.

On average, Lena had 189±12.0 and 244±11.0 OTUs in her fecal and skin samples, while the respective values for Nicole were 288±12.4 and 234±7.4 (Fig. 1, Table S1, S2). The Simpson 1-D diversity index showed little variance between the two individuals, as it averaged 0.85±0.068 and 0.87±0.026 in Lena and Nicole’s feces, respectively, while in their skin samples it averaged 0.21±0.024 and 0.93±0.021, respectively. The two individuals shared 88.8% of their fecal and 89.4% of their skin total OTUs (Fig. S1). PERMANOVA showed that the fecal and skin bacterial microbiota of the two pups were not significantly different (Table 1). However, the fecal bacterial microbiota was significantly different from the skin bacterial microbiota in both individuals (Table 1).

**Figure 1.**
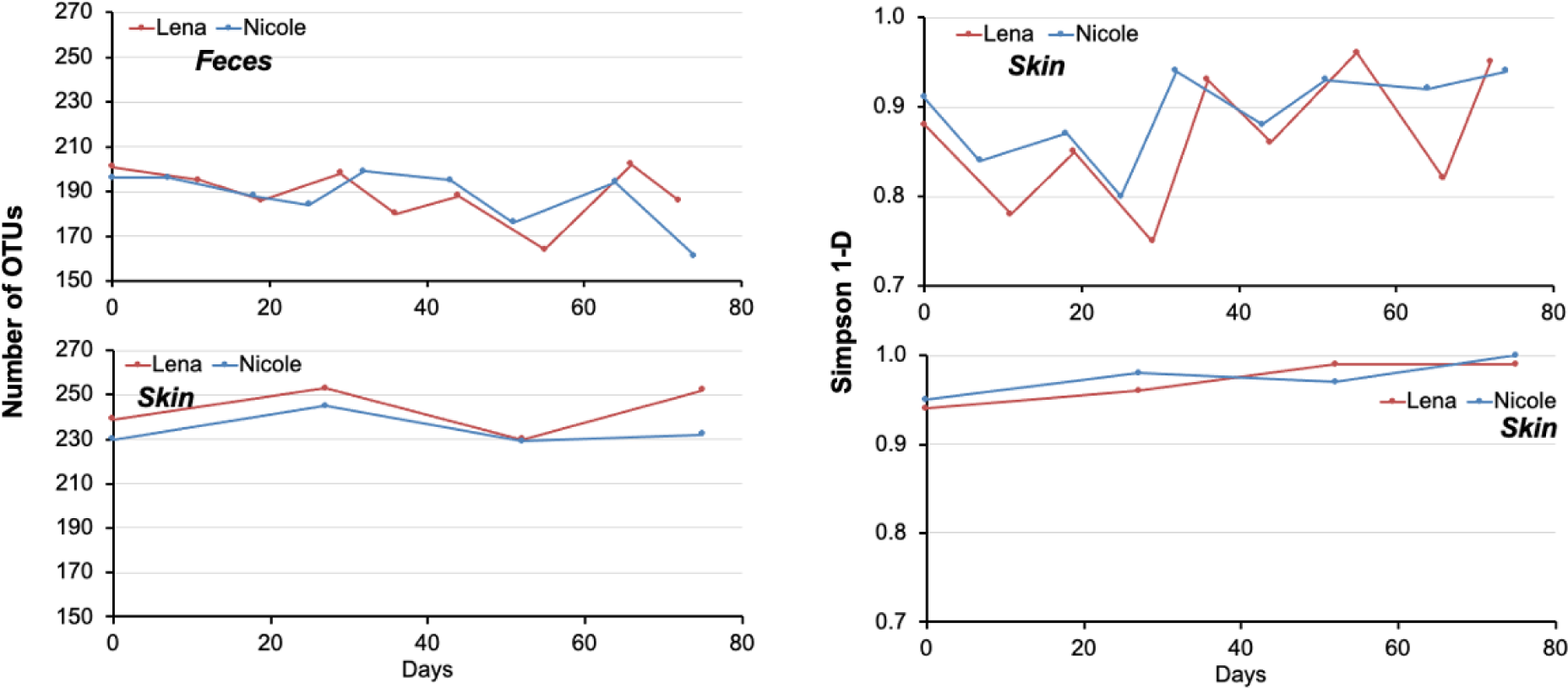
Bacterial operational taxonomic units (OTUs) richness and Simpson 1-D index in the feces and skin of two hospitalized *Monachus monachus* pups.

**Table 1.** PERMANOVA results of the fecal and skin bacterial operational taxonomic units in the feces and skin of two hospitalized *Monachus monachus* pups. Upper half are the p values and lower half of the table are the F values. Star indicates p<0.05.

|  | Lena-feces | Nicole-feces | Lena-skin | Nicole-skin |
| --- | --- | --- | --- | --- |
| Lena-feces | - | 0.689 | 0.002* | 0.001* |
| Nicole-feces | 0.568 | - | 0.002* | 0.002* |
| Lena-skin | 5.688 | 5.751 | - | 0.797 |
| Nicole-skin | 6.218 | 6.313 | 0.160 |  |

Cluster analysis of the feces bacterial microbiota, based on Bray-Curtis similarity, revealed two major groups corresponding to two time periods, with similarity of ≤40% (Fig. 2). The first group included the first three sampling dates for both pups and was dominated (≥5% relative abundance in the whole period) by OTUs affiliated with the *Shigella* (35.2% relative abundance), *Streptococcus* (17.1%), *Enterococcus* (9.7%), *Lactobacillus* (6.4%) and *Escherichia* (5.5%) genera. The second group included the rest of the samplings. During this period, the most abundant fecal bacteria OTUs were associated with the *Clostridium* (44.5% relative abundance)*, Blautia* (15.0%)*, Fusobacterium* (10.3%)*, Edwardsiella* (5.5%) and *Bacteroides* (5.3%) genera. For both individuals, statistically significant differences were found in the fecal bacterial communities between consecutive sampling points in most cases (Fig. 2). The dominant fecal OTUs (cumulative relative abundance ≥80%) of both individuals consisted of 46 OTUs (Fig. 3) with 27 of them being shared among the two individuals (Fig. S2).

**Figure 2.**
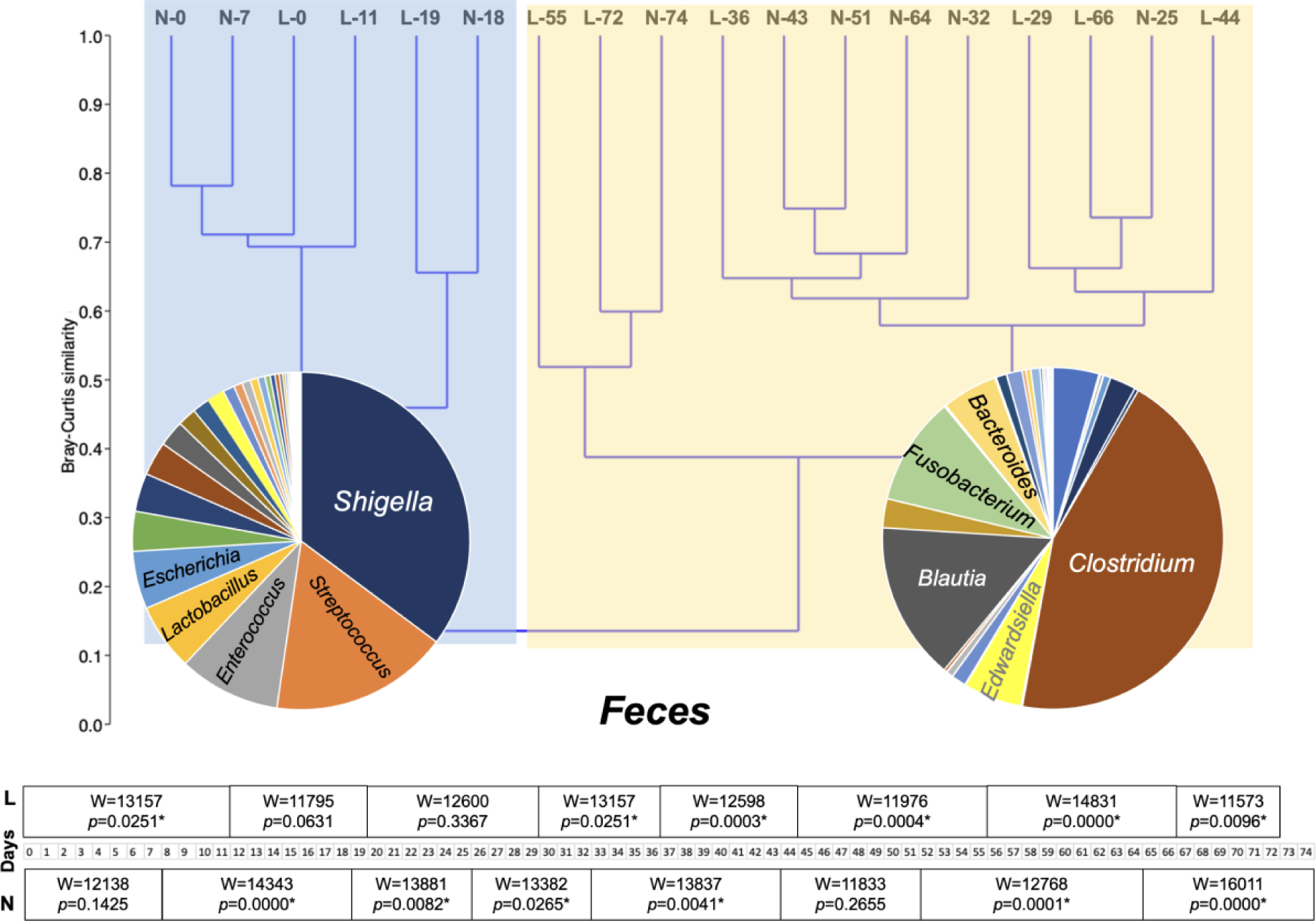
Cluster analysis of the bacterial operational taxonomic units (OTUs) abundances in the feces of two hospitalized *Monachus monachus* pups, and statistical differences (* indicates p<0.05) between consecutive sampling dates. Shaded areas represent the two major groups (see text). L: Lena, N: Nicole, L/N numbers indicate day of sampling.

**Figure 3.**
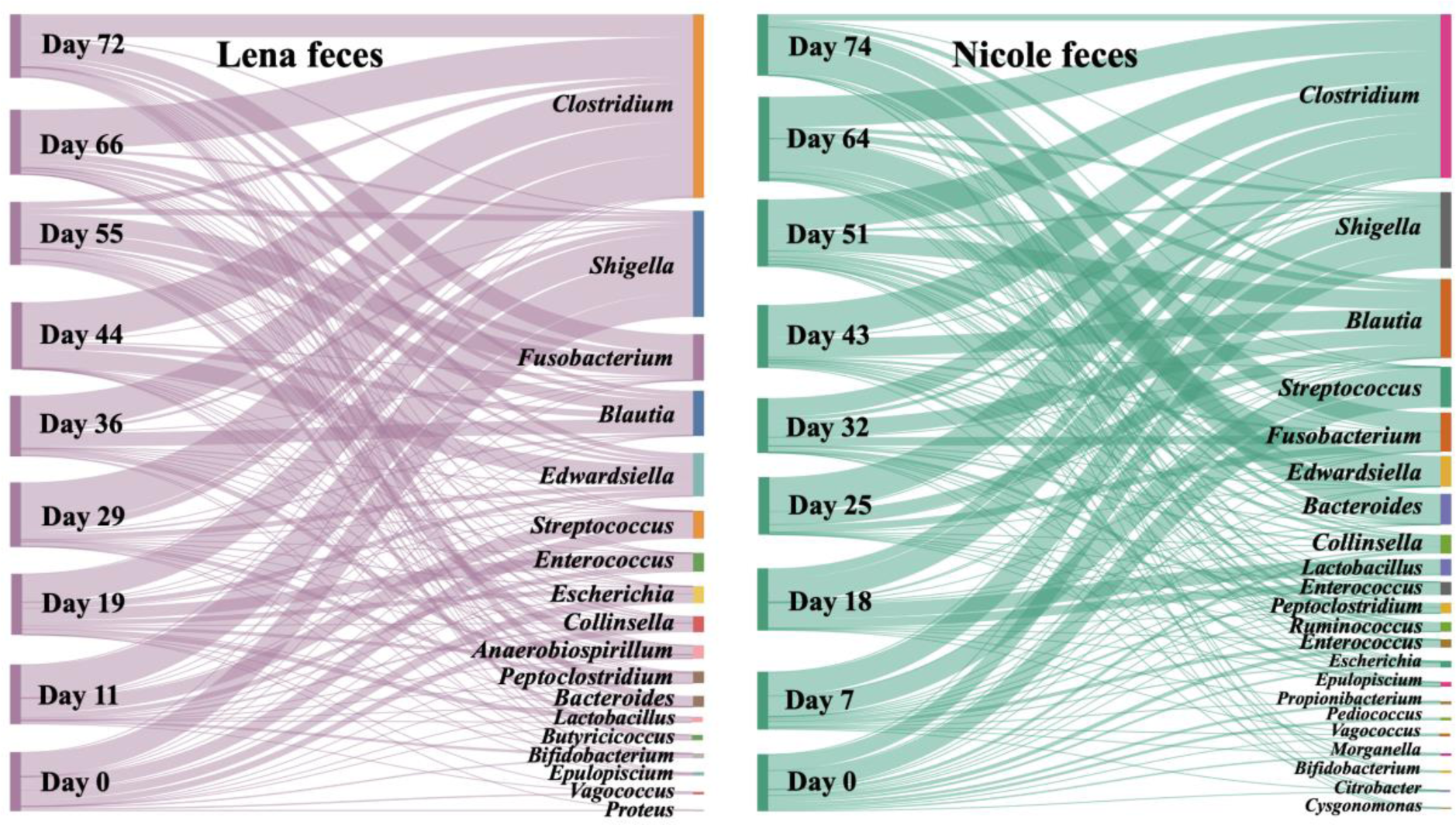
Dominant (≥80% relative abundance) bacterial genera found in the feces of two hospitalized *Monachus monachus* pups in each sampling point.

Regarding the skin bacterial microbiota, four clusters were formed with each one containing the bacterial profiles of both individuals in each sampling point (Fig. 4). In each pair, a *Psychrobacter*-related OTU dominated (44.9 – 72.2% relative abundance) while other abundant OTUs were affiliated with the *Bergeyella* (2.2 – 20.1%) and *Elizabethkingia* (0.1 – 14.8%). No statistically significant differences were found between any of the individual’s pairs, apart from day 27 (Fig. 4). A total of 41 OTUs dominated in both individuals, with 26 of them being shared among the two individuals (Fig. S2). The top dominant OTUs belonged to the *Psychrobacter*, *Bergeyella* and *Elizabethkingia* (Fig. 5). The *Psychrobacter*-like OTU over-dominated in both individuals in all sampling points, with its higher abundance in the first sampling.

**Figure 4.**
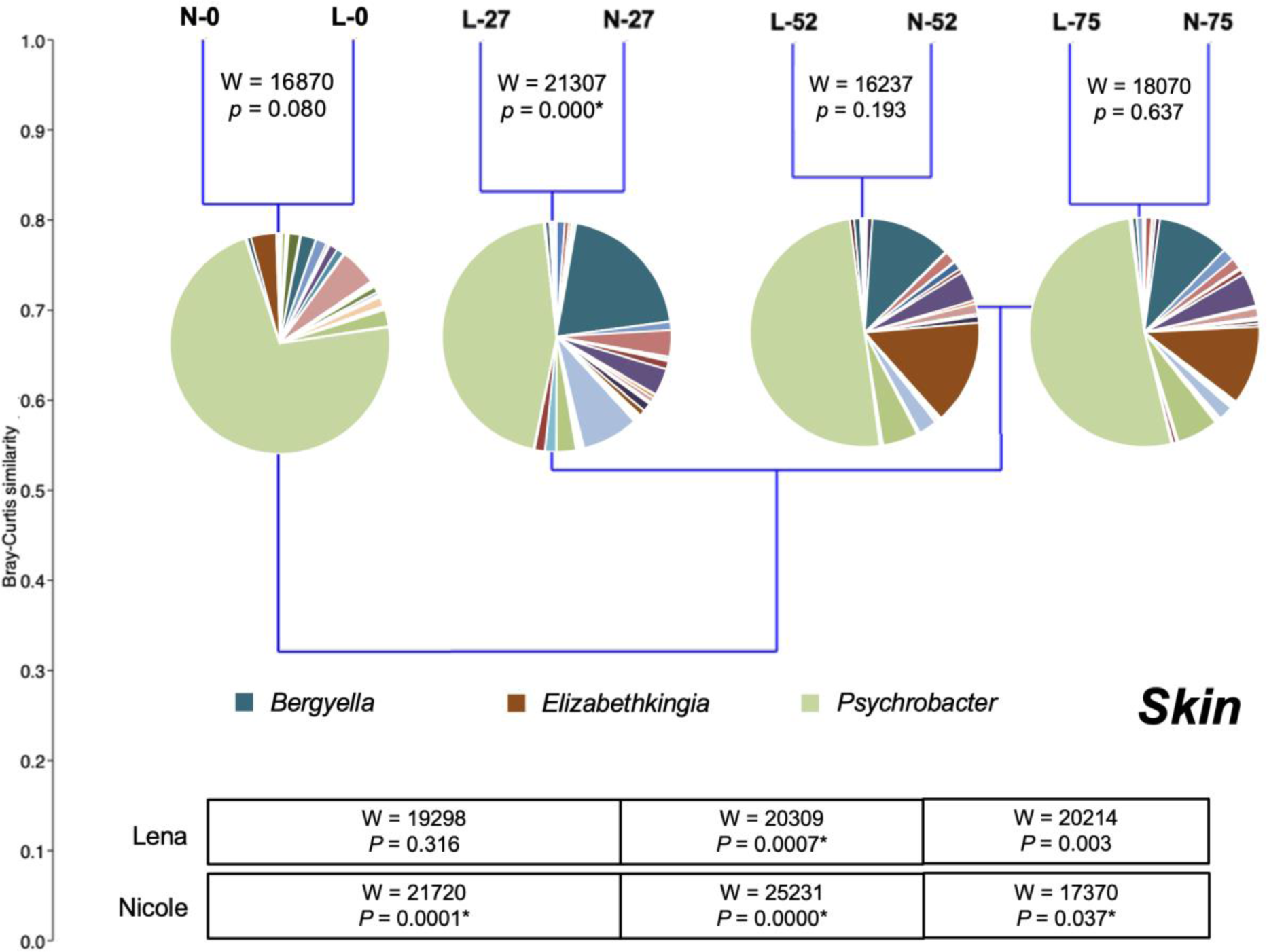
Cluster analysis of the bacterial operational taxonomic units (OTUs) abundances on the skin of two hospitalized *Monachus monachus* pups, and statistical differences (* indicates p<0.05) between consecutive sampling dates. L: Lena, N: Nicole, L/N numbers indicate day of sampling.

**Figure 5.**
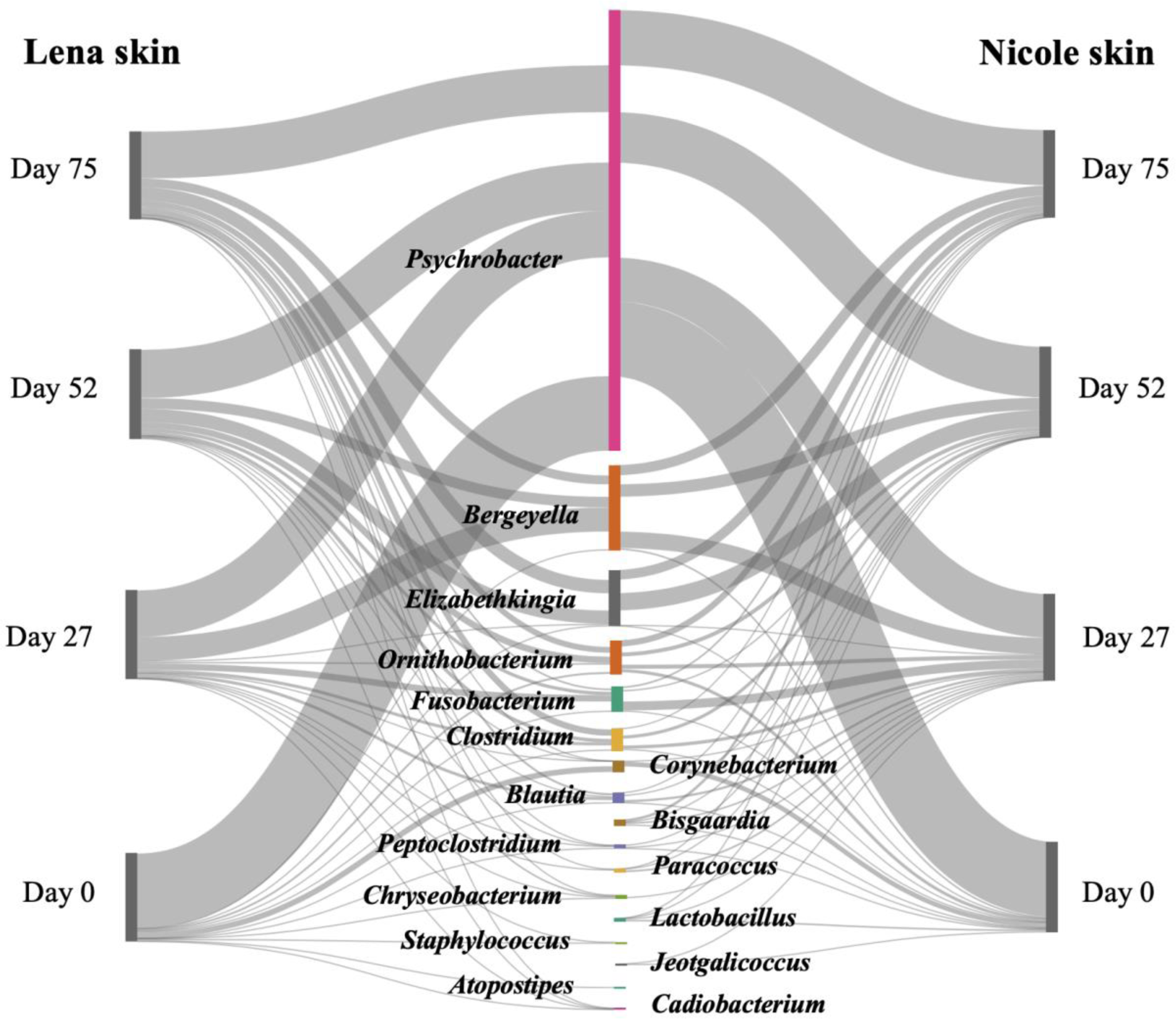
Dominant (≥80% relative abundance) bacterial genera found on the skin of two hospitalized *Monachus monachus* pups in each sampling point.

Robustness of the Lena fecal bacterial communities increased 1.8 times until day 55 and decreased 1.4 times until the last sampling (Table 2). Nicole’s fecal bacterial communities showed a continuously increased in robustness (x3.4) until the last sampling. Regarding the skin bacterial communities, robustness increased 3.0 and 2.5 times until the end of all samplings.

**Table 2.**
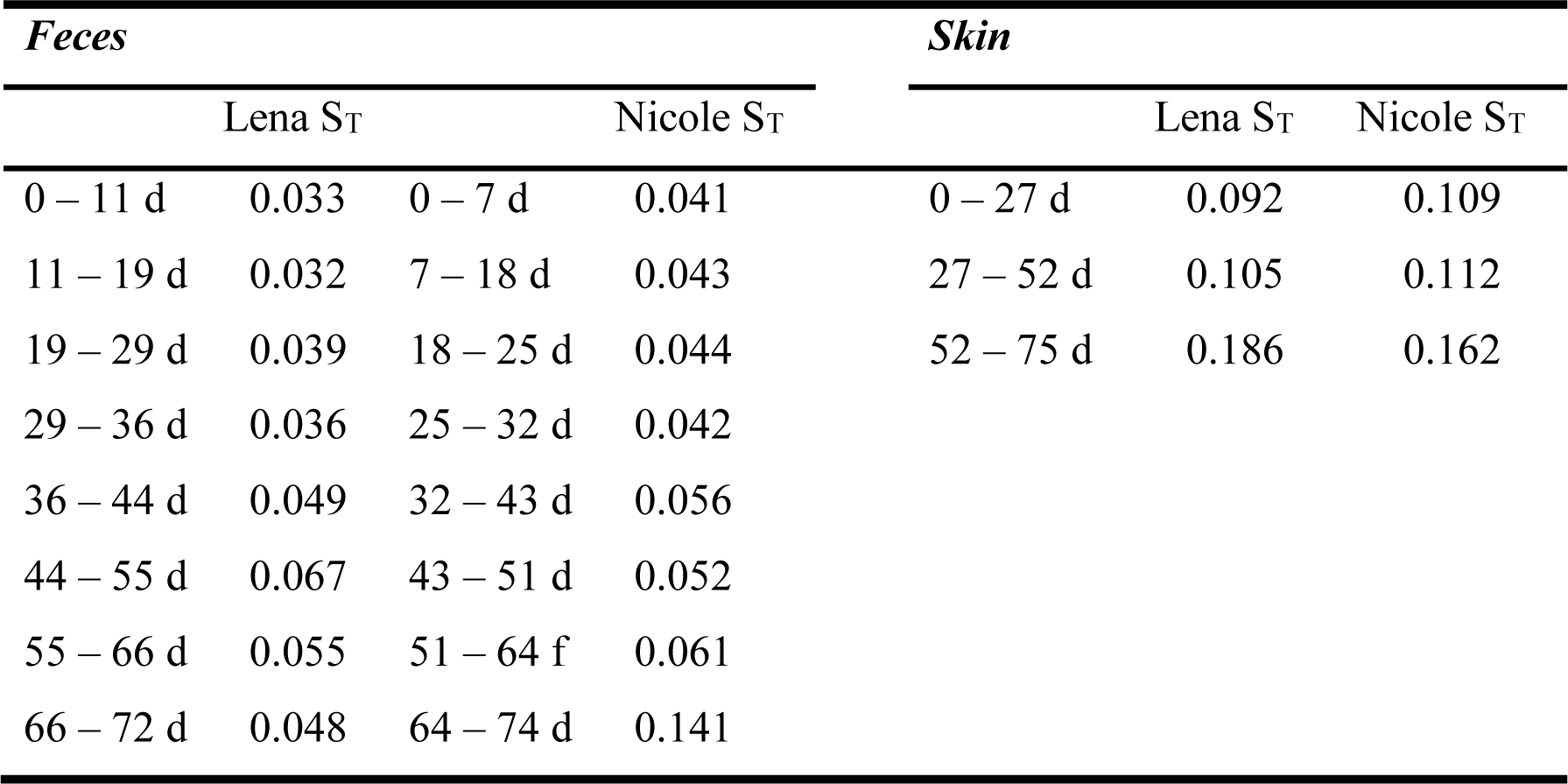
Robustness index (S_T_) of the fecal and skin bacterial communities found in two hospitalized *Monachus monachus* pups. d: day.

## DISCUSSION

Microorganisms contribute through positive and negative effects to the adaptability and fitness of their animal hosts (13) by taking part in or even regulating processes related to their host’s nutrition physiology, reproduction, development, behavior and susceptibility or resistance to infectious disease (32). Accumulating scientific literature emphasizes that host-associated microbial consortia, occurring from the animals’ skin to their gastrointestinal tracts can be informative and supporting or complementary tools to conservation practices (7, 12, 13, 33, 34). Apart from humans, the significance of microbiomes is now well accepted for both wild and domesticated or captive animals (2, 3). In addition, animal microbiomes are nowadays considered to hold important roles in the conservation of their hosts (13). For the captive animals who are in need of special care for health or rehabilitation reasons, knowing their microbiome under various environmental conditions or health statuses and how this is shaped, is becoming more important for such practices (e.g. (35, 36).

A large part of the conservation plan for the Mediterranean monk seal includes rescuing, rehabilitation, and release of stranded sick or injured adult seals or orphan seal pups by the MOm and the School of Veterinary Medicine, Aristotle University of Thessaloniki, Greece. However, to date there are no microbiome data for this species, either in natural or captive animals. Limited knowledge exists for the closely related species of the Hawaiian monk seal (*Monachus/Neomonachus schauinslandi*) regarding the presence of antibodies to specific pathogenic viruses and bacteria (29), its oral and nasal aerobic bacteria (30), and some cultivable bacterial species related to animal health (31). However, the gut and skin microbiomes are now recognized to be central for the nutrition and health of marine mammals, as for all animals (18, 37). In the present study we investigated for the first time the succession of fecal and skin bacterial communities of two rescued female Mediterranean monk seal *Monachus monachus* pups during their rehabilitation period prior to their release at sea.

The two individuals had similar bacterial profiles both in their skin and fecal microbiota, based on their statistical differences and the high number of shared OTUs, most likely due to their “common garden” (38, 39) environmental conditions, as both animals lived in the same water tank and had the same diet and health care practices. However, in each individual, their skin and fecal microbiota were different (Table 1). Such separation of the two bacterial communities has also been reported in spotted seals (40) as the gastrointestinal tract and skin select for different bacteria suggesting that their microbiota are shaped by different factors and, possibly, providing different services to their hosts.

### Fecal microbiota

The succession of the fecal bacterial microbiota was clustered in two distinct periods being ca. 40% similar to each other (Fig. 2). The first period represents, at least partially, the natural fecal microbiota of the pups, since during this period no medication or probiotics was given; the impact of the provided nutritional supplements impact is more likely to be detected in the second period. In the first period, the fecal microbiota was dominated by the *Shigella*, *Streptococcus*, *Enterococcus*, *Lactobacillus* and *Escherichia* genera. *Shigella* and *Escherichia* have been found to co-occur in the feces of captive belugas and Pacific white-sided dolphins (41), dwarf sperm whale (*Kogia breviceps*) (42) and harbor seals (43). However, it’s often considered as a disease causative agent for humans and other primates but it remains unclear if it is pathogenic to other mammals, including marine ones. *Escherichia coli* 0157 strains have been isolated from captured wild *Neomonachus schauinslandi* adult individuals, a closely related species to *M. monachus*, although the animals were not diseased, and has been hypothesized that this, along with other human-related microorganisms, are found in the seals due to their close living in human-dominated ecosystems (44). The evidence for the occurrence of antibiotic resistant bacteria in marine mammals is accumulating and concerns several of these animal species (45–48). Cultured *E. coli* prevailed in the feces of stranded-but not diseased-harbor seals (*Phoca vitulina*) individuals at admission to rehabilitation compared to wild-caught ones (49), and this is a point of concern during rehabilitation, i.e. the transfer of microorganisms from humans to the hospitalized animals. *Streptococcus* is another potentially detrimental group of bacteria but it has not been directly related to pathogenesis in marine mammals; it has been found in the oral cavity and intestine of the Yangtze finless porpoise (*Neophocaena asiaeorientalis*) (50) and sperm whales (*Physeter catodon*) (51). *Enterococcus* has been reported to be among the dominant fecal bacteria of captive or stranded but not diseased cetaceans (41, 51–55). In the present study, whether detrimental or not, the considerable decrease of *Shigella* (from 35.2% to 4.4%), *Escherichia* (from 5.5% to 0.7%), *Streptococcus* (from 17.1% to 0.1%) in the second period after the metronidazole admission, reduced any potential risks from these bacteria. *Lactobacillus*, was the last dominant group of the first period, and is well-known beneficial microorganisms of the gastrointestinal tract; indeed, Lactobacillus strains with probiotic metabolic features have been isolated from the bottlenose dolphin (*Tursiops truncatus*) (56).

In the second- and longest-period (days 29 – 66 and 25 – 64 for Lena and Nicole, respectively) of the rehabilitations process, a major turnover in the dominant OTUs was observed, dominated by the *Clostridium, Blautia, Fusobacterium, Edwardsiella* and *Bacteroides* genera. The top dominant OTUs were related to *Clostridium* with collective 44.5% relative abundance throughout this period. The genus *Clostridium* could be considered as a resident member of the pups’ microbiota, as it very frequently occurs among the most dominant bacteria found in gut and other tissues of healthy, captive, stranded and dead marine mammals ((57) and references therein). The genus contains pathogenic species, as well, with *C. perfringens* being the most frequent pathogen found in several marine mammals (58–61). In the present study, the *Clostridium*-related sequences cannot be affiliated to any of the known species of this group, but since the pups were not diseased, we hypothesize that these OTUs do not represent pathogenic members of this genus, rendering these specific bacteria as commensals, if not beneficial for the *M. monachus* pups.

To date there are no reports on the occurrence of *Blautia* in marine mammal gut or feces. This genus, which is commonly found in terrestrial mammals, has been recently suggested to hold beneficial metabolic traits for their hosts (62) and its occurrence as the second most dominant group in the second rehabilitation period, is rather desired. The genus *Fusobacterium* had 10.3% relative abundance in this period. This genus has been found in the oral cavity (50), the genitals (63) and the fecal material of other cetaceans (51, 64), as well, The phylum Fusobacteria has been found to be characteristic of marine carnivores when compared to terrestrial carnivores (28). Although it is considered to include potential pathogens to humans and animals (51, 65), in the present study its high abundance was not related to any pathogenies in the two pups. *Edwardsiella* has been associated with diseased cetaceans (66) while pathogenic species, like *E. tarda*, have been isolated from non-diseased animals (47, 67, 68). This bacterium could be a threat for the released pups, as its abundance seemed to increase towards the end of the rehabilitation period. Of similar abundance, was the genus *Bacteroides* whose abundance increased from 0.1 to 5.3% between the first and the second period. This genus has been found in the oral (50) and fecal (52, 53, 69) microbiome of cetaceans. In humans, it is considered one of the commensal bacteria that first colonize the gut after vaginal birth (70), and this could be also the case for the *M. monachus* pups.

The overall care system resulted in a very good macroscopic condition of the two pups prior to their release, and rather stable fecal bacterial communities as suggested by the robustness index, although Lena’s fecal bacterial microbiota at the last sampling was slightly less stable compared to the rest of the samplings. The antibiotic’s impact on the fecal microbiota eliminated the potentially detrimental *Shigella* dominance on the first period. The probiotic’s impact on the fecal microbiota of the pups could not be evaluated in the present study as it was administered on day 72 and day 64 for Lena and Nicole, respectively. One of the two pups was photographed (identified by the presence of a marker tag) in a healthy state circa 6 months after release in a regularly monitored seal cave.

### Skin

The skin bacterial microbiota showed a converging pattern from the beginning to the end of the rehabilitation period, as a result of adaption to the artificial environment of the rehabilitation water tank. No statistical differences between the two pups were observed except in day 27. These final skin bacterial communities, prior to the release of the animals, seem to be more stable than the initial ones, as assessed by the robustness index. The observed statistically significant differences in all but one case, between consecutive samplings for each pup Fig. (4), are attributed to the changes in relative abundance of the same OTUs as the overlap in OTUs occurrence was high (Fig. S1).

The three dominant (Fig. 4, 5) genera *Psychrobacter, Elizabethkingia* and *Bergeyella* showed different patterns during the rehabilitation. Despite that *Psychrobacter*-related OTUs dominated in each sampling, their relative abundance decreased gradually during the rehabilitations period. This genus is common on the skin of marine mammals (50, 71–73), reflecting its ubiquity in the marine environment (74, 75). Although it remains unclear if some of its species could be true pathogens, an extensive comparative genomics study concluded that the genus’s strains belong to either the flexible ecotype, which can grow at warm temperatures and so it can colonize mammalian skin and other tissues, or the restricted ecotype, i.e., the pure psychrophiles, free-living and generalist strains found in the world ocean. Both ecotypes, have a pathobiont evolutionary origin, whose virulence was lost or weakened via genome reduction (76). Its dominance from the pups’ transition from the natural marine to the artificial environment of the rehabilitation water tanks, is in accordance with its high adaptability (74, 76).

In our study, the *Elizabethkingia*-related bacteria were temporarily favored during the rehabilitation period, but at the last sampling they decreased to >0.1%. Despite that to date no reports exist on the occurrence of *Elizabethkingia* in marine mammals, these microorganisms are of special interest as some of its species have been fish-associated and reported as pathogenic or spoilage microorganisms (77). Moreover, infections of *E. meningoseptica* are considered serious and dangerous in humans (78), cows (79) and frogs (80). It is possible that the provided metronidazole in the two seal pups, controlled the uprising of the *Elizabethkingia*-related bacteria towards the last half of their rehabilitation period.

*Bergeyella*-related bacteria is another group of skin-associated microorganisms which was favored under the rehabilitation conditions. Its dominance increased between the first and the last sampling, possibly not affected by the provided antibiotic. Although there are no reports on *Bergeyella* in seals, members of this genus have been found with higher abundance in bottlenose dolphins (*Tursiops truncatus*) calves compared to adult males (81). Although in the present study the species identification is of limited security due to the inherent decrease predictability of the short sequence lengths of high-throughput sequencing, some species of this genus could pose potential risk to the pups, such as *Bergeyella zoohelcum* which is a pathogen of the upper respiratory tract of dogs, cats and other mammals (82).

For the successful reintroduction processes of protected animal species, such as the Mediterranean monk seal (*Monachus monachus*), the final aim of any rehabilitation process is to have healthy animals prior to their release in the wild. Both detrimental and beneficial microorganisms associated with these animals are of central importance for the reintroduction to the wild, with housing and care conditions being at the frontline (83). The present study investigated for the first time the skin and fecal microbiota succession of two rescued female *M. monachus*) pups during their rehabilitation period. It revealed very low individual variability in both skin and fecal microbiota and some dominant bacterial genera which have been reported for the first time in *M. monachus*, or even marine mammals in general. The forecasting power of microbiomes (84) along with its now fast advancing microbiome engineering (85, 86) even for conservation (13) and halting biodiversity loss (7) opens the way for moving from observational to interventional and functional microbiome research for the protected Mediterranean monk seal.

## MATERIALS AND METHODS

### Rehabilitation

Two female *M. monachus* pups, named as Lena and Nicole, were admitted in the MOm-Monk Seal Rehabilitation Centre, Attiki, Greece, on 24 Sep. and 10 Oct. 2019, respectively. The pups were found stranded in the Eastern Peloponnese (Lena) and North-East Euboea (Nicole) on 23/09/2019 (Lena) and 09/10/2019 (Nicole) and weighted 15.9 kg, (Lena) and 16.5 kg (Nicole). The estimated pups’ age was 7 and 20 days for Lena and Nicole, respectively. The animals spent 130 (Lena) and 113 (Nicole) days under rehabilitation conditions before being released back in the wild. The seal holding area included a dry platform (ca. 9m^2^) and a sea water tank (ca. 10m^3^). Recirculating tank water temperature was kept at 17±2°C. The sea water was brought in from the nearby coast of Artemida by tanker truck and was renewed on average every 6 days. This water was treated continuously by a protein skimmer and every 2-3 days with controlled chlorine doses. The tank water was monitored visually daily and the origin water was tested monthly. The pups were fed with *Scomber scomber,* which was provided as fish porridge (up to days 49 and 40 for Lena and Nicole, respectively), combination of fish porridge and whole fish (between days 49-79 and 40-61 for Lena and Nicole, respectively) and for the rest of the rehabilitation time as whole fish. Feeding frequency ranged from one to six times per day, depending on the growth stage of the pups. On their release date, the pups were clinically healthy and weighted 57.4 kg (Lena) and 55.0 kg (Nicole).

To prevent dehydration, electrolytes solution of Almora sachets PLUS (Elpen, Greece) was provided from start alone the first 48 hours and then and in fish porridge to day 79 (Lena) and 61 (Nicole). Boiled Quaker oats extract was provided for the first 8 (Lena) and 5 (Nicole) days. One Aquavits tablet (International Zoo Veterinary Group, UK) was given once daily between days 6 and 109 for Lena and days 12 and 91 for Nicole. PetCal (Zoetis, USA) was used as a supplement of phosphorous, calcium, and vitamin D3, given as 1.5-5 tablets/d from day 6 (Lena) and day 12 (Nicole) until the end of their rehabilitation period. Lena was given daily dosages of 4-20 ml of SalmoPet salmon oil (MarinPet, Norway) in days 5-17 and 30-41 and Nicole 5-15 ml in days 16-23. Finally, one sachet of the probiotics supplement PURINA PRO PLAN Canine FortiFlora (Purina, USA) was given to Lena daily between days 72-82 and then every four days up to day 125. Nicole got the same probiotics supplement with the same dosage between days 53-61 and after that she was receiving a single sachet every three or four days until day 111.

Metronidazole was given as prophylaxis to both pups. Lena received daily 5mg/kg body weight of metronidazole between days 39-70. The respective daily dosage for Nicole was the same between days 20-50. Nicole was treated twice a day with azithromycin eye drops (Azyter; Laboratoires Thea, France) from days 6-17 and days 29-30.

### Molecular analyses and data processing

A total of nine individual fecal and four skin swab samples were collected and analyzed for each individual during their rehabilitations period. Our first sampling (day 0) corresponds to the second day of the pups in the rehabilitation center. Fecal samples were collected immediately upon defecation from each individual. Pre-sterilized cotton swab scrapings were retrieved along the sides of each individual. All samples were immediately frozen at −20°C and then at −80°C. Bulk DNA was extracted from ca. 0.3 g of fecal material or the whole cotton swab using the NuceloSpin Soil DNA extraction kit (Machery-Nagel, Germany) according to the manufacturer’s guidelines.

For the PCR amplification of the V3–V4 regions of the bacterial 16S rRNA gene from the bulk extracted DNA, we used the primer pair S-D-Bact-0341-b-S-17 and S-D-Bact-115 0785-a-A-21 (87). The amplified sequences were sequenced on a MiSeq Illumina instrument (2×300 bp) at the MRDNA Ltd. (Shallowater, TX, USA) sequencing facilities. Unprocessed DNA sequences are available in the Sequence Read Archive (https://www.ncbi.nlm.nih.gov/sra/) under BioSample SAMN32536541 of the BioProject PRJNA917309. All processing of the raw 16S rRNA gene sequences was performed by using the MOTHUR standard operating procedure (v.1.46.1) (88, 89). The resulting operational taxonomic units (OTUs) were grouped as identical at 97% cut-off similarity level and were classified with the SILVA database release 138 (90, 91). For those OTUs which were designated as “unaffiliated”, their closest relatives were found by using Nucleotide Blast (http://blast.ncbi.nlm.nih.gov).

Data and statistical analysis and graphic illustrations were performed using Palaeontological STudies (PAST) software (92) and the vegan package (93) in R Studio platform Version 1.1.419 (94) with 3.4.3 R version. We applied cluster analysis based on the unweighted pair group method with arithmetic mean Bray-Curtis similarity. Permutational multivariate analysis of variance (PERMANOVA) was used to detect differences between the fecal and skin bacterial microbiota of the two pups and the Wilcoxon test was applied to detect the microbiota differences between sequential samplings. The robustness of the fecal and skin bacterial microbiota to temporal stability and resistance were investigated in this study. Robustness is an index which assesses the degree of a community’s structural constancy over time (95), its ability to resist change following perturbation (96, 97), and its resilience, i.e., its ability to return to an initial structure following perturbation (98).

## Supporting information

Supplementary material contains two tables and two figures

## ACKNOWLEDGEMENTS

We would like to thank the Attica Zoological Park for the logistical and financial support of the two orphan monk seal pups. The rehabilitation of the two pups was conducted following strict protocols and with all necessary permits from the national relevant authorities of Greece.

## CONFLICT OF INTEREST

The authors declare no conflict of interest.

## AUTHOR CONTRIBUTIONS

Kon. K. and A.D. conceived the ideas, designed methodology and led the writing of the manuscript; A.D., A.M. and Kon. K. performed microbiota and bioinformatics and data analysis; A.D., Kim. K., A.K., E.T. and P.D. performed all sampling and animal care during rehabilitation. A.K. was responsible for the veterinary care of the animals. All authors contributed critically to the drafts and gave final approval for publication.

## Notes

### Competing Interest Statement

The authors have declared no competing interest.

