## Supplementary material contains two tables and two figures for "Fecal and skin microbiota of two rescued Mediterranean monk seal pups during rehabilitation"

**Table S1**. Sequencing results and alpha diversity indices of the fecal bacterial communities of two rehabilitated *Monachus monachus* pups (named Lena and Nicole). OTU: operational taxonomic unit.

|  | ***Lena*** | | | | | | | | |
| --- | --- | --- | --- | --- | --- | --- | --- | --- | --- |
| *Sampling day* | 0 | 11 | 19 | 29 | 36 | 44 | 55 | 66 | 72 |
| *Reads* | 31458 | 31585 | 29817 | 31462 | 29775 | 31268 | 29772 | 32069 | 31671 |
| *No. of OTUs* | 201 | 195 | 186 | 198 | 180 | 188 | 164 | 202 | 186 |
| *Simpson 1-D* | 0.83 | 0.73 | 0.80 | 0.70 | 0.88 | 0.81 | 0.91 | 0.77 | 0.90 |
| *Shannon H’* | 2.53 | 2.13 | 2.37 | 2.19 | 2.97 | 2.42 | 2.87 | 2.53 | 3.06 |
| *Chao-1* | 239.9 | 257.6 | 256.7 | 245.5 | 209.5 | 270.6 | 216.6 | 253.8 | 233.3 |
|  | ***Nicole*** | | | | | | | | |
| *Sampling day* | 0 | 7 | 18 | 25 | 32 | 43 | 51 | 64 | 74 |
| *Reads* | 30404 | 30824 | 28192 | 31004 | 30772 | 29702 | 30533 | 29320 | 31154 |
| *No. of OTUs* | 196 | 196 | 188 | 184 | 199 | 195 | 176 | 194 | 161 |
| *Simpson 1-D* | 0.86 | 0.79 | 0.82 | 0.75 | 0.89 | 0.83 | 0.88 | 0.87 | 0.89 |
| *Shannon H’* | 2.77 | 2.44 | 2.25 | 2.38 | 2.95 | 2.59 | 2.75 | 3.05 | 2.81 |
| *Chao-1* | 255.4 | 257.9 | 241 | 242 | 232.4 | 238.4 | 230.1 | 233 | 218.4 |

**Table S2**. Sequencing results and alpha diversity indices of skin bacterial communities of two rehabilitated *Monachus monachus* pups (named Lena and Nicole). OTU: operational taxonomic unit.

|  | ***Lena*** | | | |
| --- | --- | --- | --- | --- |
| *Sampling day* | 0 | 27 | 52 | 75 |
| *Reads* | 31998 | 32064 | 32329 | 32102 |
| *No. of OTUs* | 239 | 253 | 230 | 252 |
| *Simpson 1-D* | 0.89 | 0.91 | 0.94 | 0.94 |
| *Shannon H’* | 3.09 | 3.26 | 3.34 | 3.51 |
| *Evenness J'* | 0.092 | 0.102 | 0.123 | 0.132 |
| *Chao-1* | 272.8 | 288.5 | 295.1 | 287.7 |
|  | ***Nicole*** | | | |
| *Sampling day* | 0 | 27 | 52 | 75 |
| *Reads* | 31963 | 31718 | 32534 | 31901 |
| *No. of OTUs* | 230 | 245 | 229 | 232 |
| *Simpson 1-D* | 0.90 | 0.93 | 0.92 | 0.95 |
| *Shannon H’* | 3.07 | 3.56 | 3.14 | 3.52 |
| *Evenness J'* | 0.093 | 0.143 | 0.101 | 0.145 |
| *Chao-1* | 283.5 | 280.8 | 309 | 293.9 |

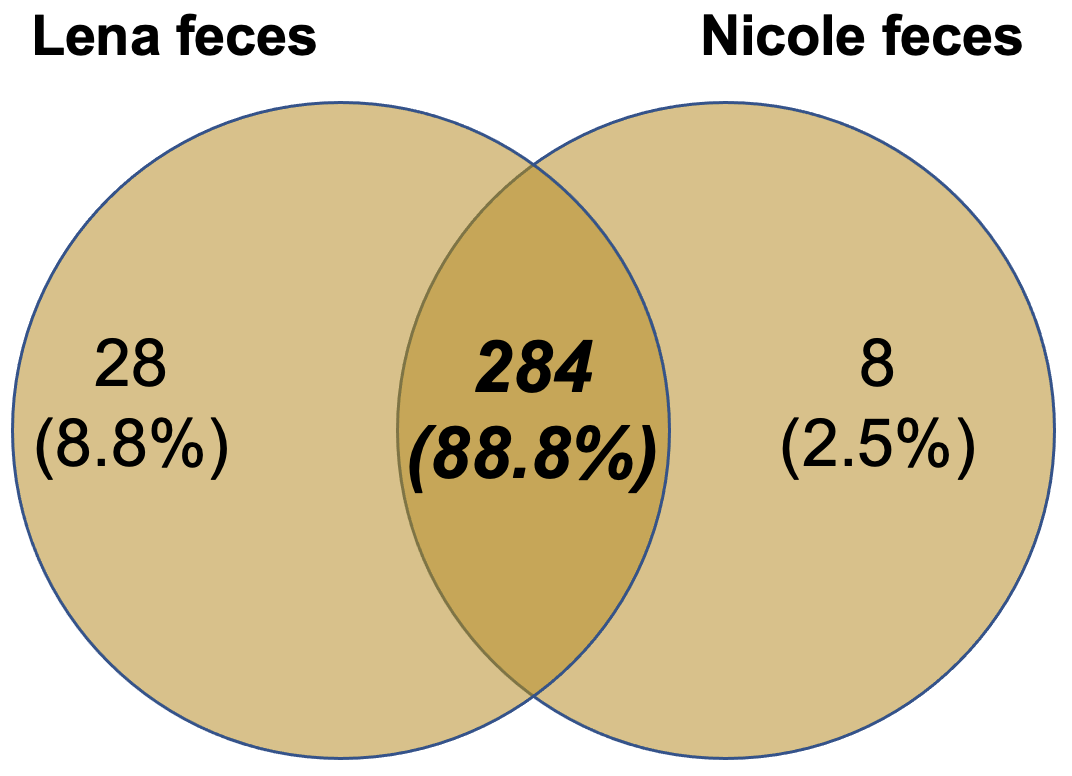

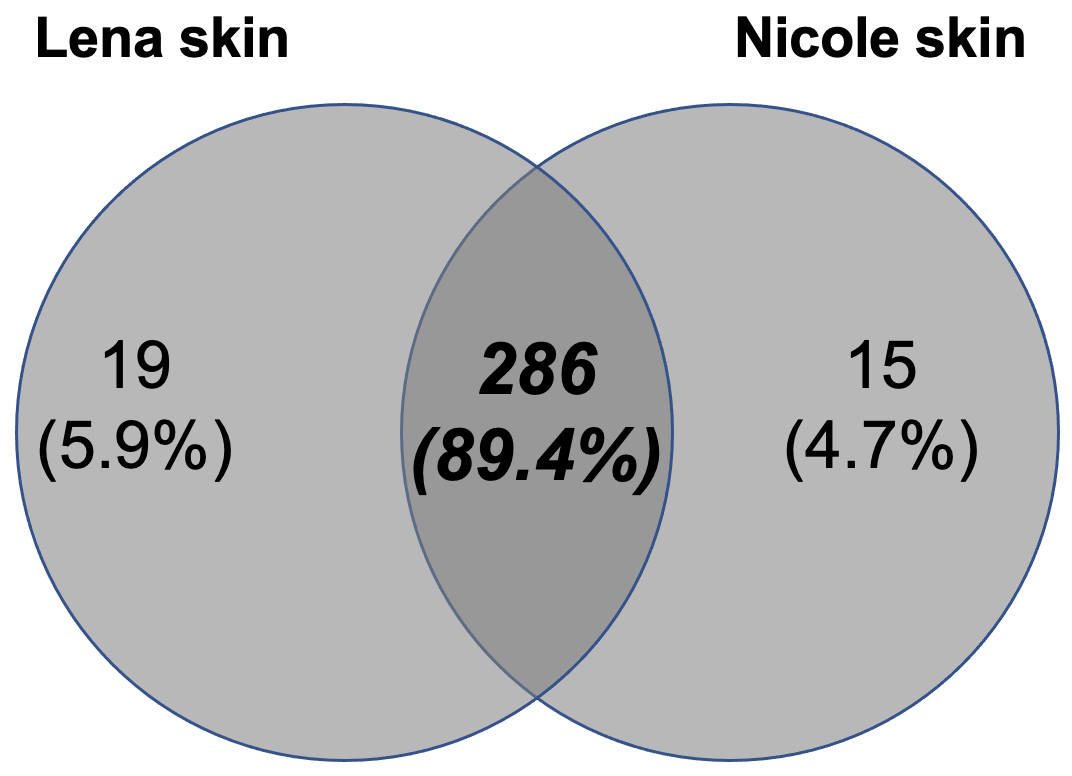

**Figure S1**. Number of shared and unique bacterial operational taxonomic units (OTUs) in the feces and skin of two rehabilitated *Monachus monachus* pups.

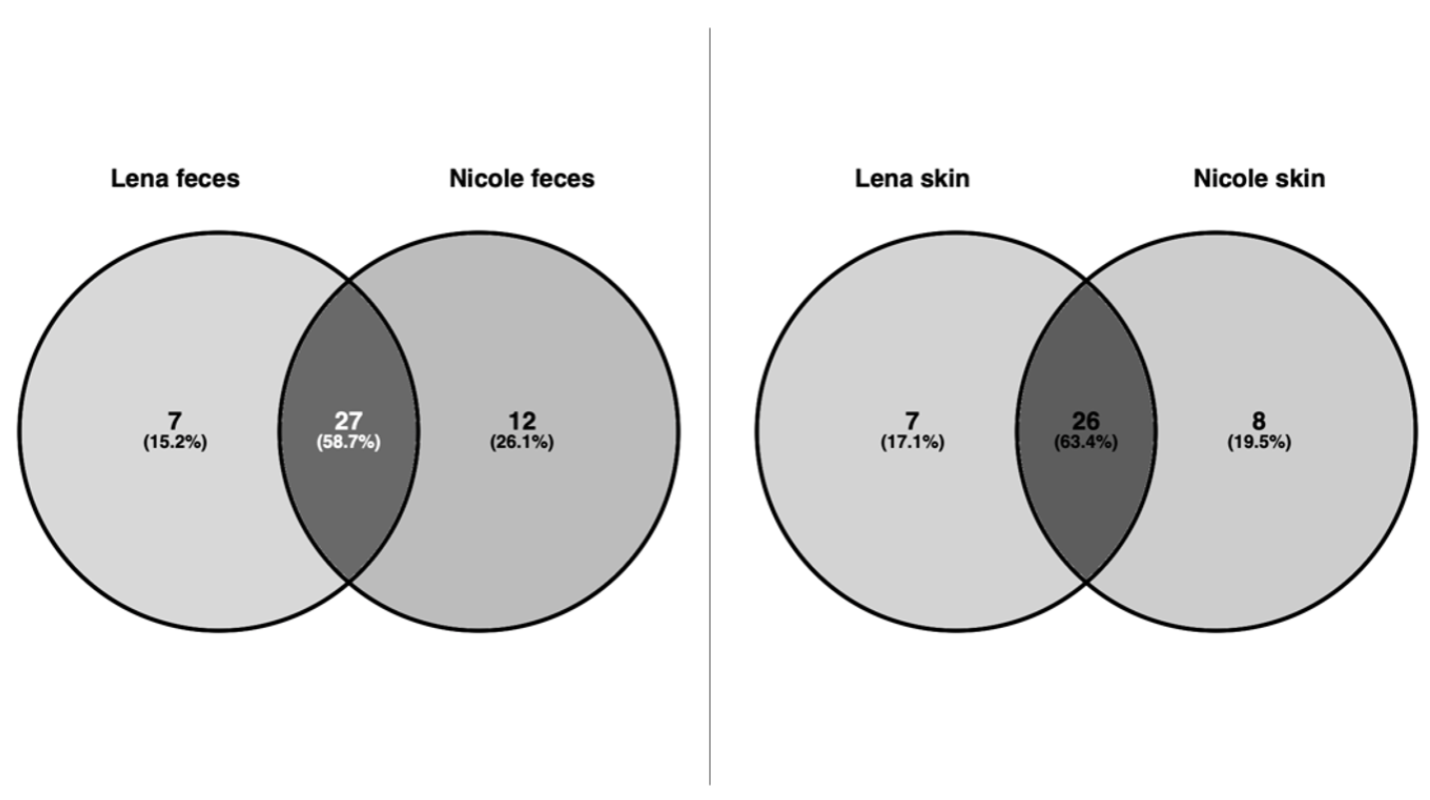

**Figure S2**. Number of shared and unique dominant (≥80% relative abundance) operational taxonomic units in the feces and skin of the two rehabilitated *Monachus monachus* pups.
